# Muscle-3D: scalable multiple protein structure alignment

**DOI:** 10.1101/2024.10.26.620413

**Authors:** Robert C. Edgar, Igor Tolstoy

## Abstract

Protein multiple alignment is an essential step in many bioinformatics analysis such as phylogenetic tree estimation, HMM construction and critical residue identification. Structure is conserved between distantly-related proteins where amino acid similarity is weak or undetectable, suggesting that structure-informed sequence alignments might offer advantages over alignments constructed from amino acid sequences alone. The advent of the AI folding era has unleashed millions of high-quality predicted structures, motivating the development and assessment of scalable multiple structure alignment (MStA) methods. Here, we describe Muscle-3D, a new MStA algorithm combining a rich sequence representation of structure context, the Reseek “mega-alphabet”, with state-of-the art alignment techniques from Muscle5 including a posterior decoding pair-HMM, consistency transformation, iterative refinement and ensemble construction. We show that Muscle-3D readily scales to thousands of structures. Comparative validation on several benchmark datasets using different quality metrics shows Muscle-3D to be among the higher-scoring methods, but we find that algorithm rankings from different metrics disagree despite low P-values according to the Wilcoxon rank-sum test. We suggest that these conflicts arise from the inherently fuzzy nature of structural alignment, and argue that a universal standard of MStA accuracy is not possible in principle. We describe contact map profiles for visualizing variation in inter-residue distances, and introduce a novel measure of local conformation similarity, LDDT-muw.

Muscle-3D software is available at https://github.com/rcedgar/muscle.

## Introduction

A multiple sequence alignment (MSA) arranges protein sequences into a rectangular matrix with the goal that residues in a given column should be equivalent. Desired criteria for equivalence may be homology (related by descent from the same position in an ancestral sequence), or superposition in a rigid local structural alignment, or common functional role. Although these criteria agree for closely related proteins, sequence, structure and function diverge over evolutionary time such that different criteria may result in different alignments. In particular, sequence similarity and secondary structure may be shifted relative to each other by one or more residues (Cline et al., 2002; Edgar, 2010). A multiple structure alignment (MStA) quantifies similarity between two or more protein structures, typically based on the coordinates of backbone *C_α_* atoms only. For closely-related proteins, the alignment may be specified by a translation and rotation of the coordinates of one structure so that similar segments overlap in three-dimensional space, enabling visualization using a tool such as Pymol (DeLano et al., 2002).

More commonly, a MStA is specified as an amino acid (a.a.) MSA where columns represent residues inferred to be structurally equivalent. Internally, an MStA algorithm may construct local superpositions of structure segments from which the MSA is inferred, but these superpositions are often not reported and are rarely used in practical applications. Thus, from a user’s perspective, an MStA is often nothing more than an MSA whose structures are available for inspection, and conversely an MSA needs only a corresponding set of structures to be considered as a viable MStA. In our terminology, MStA refers exclusively to an a.a. multiple alignment in conventional matrix format which was constructed with the help of structural information.

## Protein of Theseus

The paradox of the Ship of Theseus (Scaltsas, 1980) asks whether a ship is the same object after having some or all of its original components replaced over time. The same question can be asked of distantly-related proteins. If mutations are exclusively substitutions, then the line of descent of each residue is maintained. But when overlapping insertions and deletions occur, some residue lineages may go extinct while preserving recognizably similar secondary structure elements. In this scenario, it may be meaningful to identify a secondary structure, say an alpha helix, as homologous between a pair of structures, while individual residues in that helix cannot be traced back to a common ancestral residue, even if complete knowledge of evolutionary history is available. Therefore, a residue found at apparently equivalent positions in the secondary structure (say, the third residue in a helix) may be homoplastic (i.e., playing the same role in both proteins) without necessarily being homologous. When the length or conformation of a secondary structure element varies, it is no longer possible to unambiguously identify the “same position”; correspondences between residues become somewhat arbitrary so that different people and different algorithms might reasonably disagree. In the ship metaphor, components are not necessarily replaced in a one-to-one fashion—a third mast may be added beside the original two masts, or the dining area divided into two cabins (secondary structure elements may be inserted, deleted or change conformation such that e.g. a helix becomes a loop). When one-to-one correspondence is lost, equivalence of a larger object may be preserved while identity of some of its component parts is not.

These considerations show that for distantly related proteins, conventional alignments presented as a rectangular matrix are fuzzy—in general, there is no “correct” alignment for many residues, even in principle. If correctness is not well-defined, accuracy cannot be measured and assessment of MStAs by comparison with putative gold-standard reference alignments is potentially biased towards the chosen criteria built into those references, raising a challenge for algorithm validation.

## MStA algorithms

The conceptually simplest approach to aligning two protein structures is to seek a rigid-body transformation of one backbone which minimizes distances between corresponding *C_α_* atoms (Kabsch, 1976; Umeyama, 1991). With distantly-related proteins, three problems arise: (1) similarity may be limited to a sub-chain of one or both structures (e.g., a domain which is flanked by unrelated domains) so that local rather than global alignment is required, (2) some regions may “flex” relative to others over evolutionary time so that somewhat different transformations should be applied to different segments, and (3) the number of *C_α_* atoms varies due to insertion and deletion mutations causing pair-wise correspondences to become unclear (for example, choosing the closest *C_α_* may result in asymmetrical or many-one correspondences). For multiple alignment of three or more proteins, these issues are correspondingly more challenging. Published algorithms based on *C_α_* coordinates include Mustang (Konagurthu et al., 2006), Matt (Menke et al., 2008) and USalign (Zhang et al., 2022). An alternative approach is to represent a structure as a sequence of letters, one for each *C_α_*, and adapt MSA algorithms such as progressive alignment (Feng and Doolittle, 1987) to MStA. This approach was recently implemented in Foldmason (Gilchrist et al., 2024), building on the 20-letter 3Di structure alphabet introduced by Foldseek (Van Kempen et al., 2024). Here, we describe Muscle-3D, a new MStA algorithm combining a rich structure alphabet with hidden Markov model (HMM) alignment, consistency transformations and iterative refinement.

## Viral RNA-dependent RNA polymerase

To illustrate analyses enabled by Muscle-3D, we use viral RNA-dependent RNA polymerase (RdRp) which is widely used in RNA virus metagenomics (see e.g. (Edgar et al., 2022)). RdRp has well-conserved motifs which are conventionally named A, B, C, D, E, F1 and F2 (Te Velthuis, 2014). Some motifs have little detectable sequence similarity in novel viruses, raising challenges for identifying and classifying diverged RdRps and RdRp-like sequences in metagenomic assemblies (Edgar, 2023).

## Methods

### Muscle-3D

Muscle-3D is a variant of Muscle5 (Edgar, 2022), a state-of-the-art MSA algorithm, in which the Re- seek “Mega” alphabet (Edgar, 2024) is used to represent structure in place of the standard a.a. alphabet. Muscle5 exploits several techniques to achieve the highest accuracy reported to date on MSA benchmarks, including posterior-decoding (maximum expected accuracy) alignment by a pair HMM, (Do et al., 2005), consistency transformations (Notredame et al., 2000) to reconcile transitive pair-wise alignments which may not agree, and iterative refinement (Gotoh, 1996). Divide-and-conquer strategies (Smith, 1985) are used to enable scaling to thousands of sequences. Ensembles of alternative MSAs can be generated to assess robustness of the alignment and of downstream inferences such as phylogenetic trees.

### MSA benchmarks

The *de facto* standard benchmark for MSA methods is Balibase (Thompson et al., 2005), a collection of reference MSAs with annotated “core blocks” which are considered to be well-aligned. Most core blocks columns have no gaps; letters outside core blocks are not aligned. At the time Balibase was constructed, solved structures were available for roughly one third of the sequences. These structures were used to inform the identification of core blocks; other proteins were necessarily aligned based only on their a.a. sequences. Balibase is divided into full-length (BB) and trimmed (BBS) subsets. BB proteins have a common domain surrounded by flanking domains which are often very different and thus clearly not homologous. The BBS subset is derived from BB by trimming to the common domain to obtain globally alignable sequences; reference alignments for BBS and BB are identical. Accuracy of a tested MSA is assessed by determining *TC*, the fraction of core block columns in the corresponding reference MSA which are fully reproduced across all sequences. Accuracy of an algorithm can be summarized as its mean *TC*, and the Wilcoxon rank-sum test (Fay and Proschan, 2010) can be used to determine whether one algorithm is significantly better than another. Homstrad (Mizuguchi et al., 1998) is another collection of reference MSAs which also has been used to benchmark MSA and MStA algorithms including Clustal-omega (Sievers and Higgins, 2018) and Foldmason. Balibase MSAs have a minimum of four proteins with a mean of 29; by contrast, most Homstrad alignments (630/1032) are pair-wise with only 233 (22%) having four or more. All Homstrad proteins are globally alignable, i.e. have the same domain organization, and all had solved structures at the time its reference alignments were constructed. As with Balibase, the alignments were manually adjusted according to criteria which are not specified in detail and are necessarily somewhat subjective. Homstrad reference alignments include all letters from all proteins, while Balibase references are specified only in core blocks. Thus, Homstrad emphasizes pair-wise alignments of globally alignable proteins, while the Balibase BB subset comprises more challenging cases requiring alignment of many proteins where homologous segments are surrounded by highly diverged and/or unrelated sequence.

### Balibase3D

We identified predicted structures in the AlphaFold Structure Database (AFDB) (Varadi et al., 2022) corresponding to Balibase sequences lacking a solved structure in the Protein Data Bank (PDB) (Berman et al., 2003). Starting from the BB subset, we discarded sequences for which no structure was found in PDB or AFDB, and then discarded accessions with less than four structures. This gave 197 sets of at least four structures each having a common domain; we call this collection Balibase3D.

### Balifam3D

Balifam (Edgar, 2022) is an MSA benchmark constructed by expanding Balibase with homologs identified via in PFAM HMMs (Bateman et al., 2004). Similarly to Balibase, we identified PDB and AFDB structures for sequences in Balifam, calling the resulting collection Balifam3D. Following the original Balifam, Balifam3D has three subsets with ~100, ~1, 000 and ~10, 000 structures per set, here called Balifam3D-100, Balifam3D-1k and Balifam3D-10K respectively. Balifam3D-100 has 59 sets.

To keep data sizes and software runtimes for comparative benchmarking within our budget limitations, Balifam3D-1k was reduced to ten sets and Balifam3D-10k four.

### Reference-free MStA quality metrics

The “Protein of Theseus” paradox implies that a universal gold standard for MStA correctness is not possible in principle. Alternatively, an MStA can be evaluated by an intrinsic quality metric calculated from the alignment plus structures. The DALI *Z* score (Holm and Sander, 1993) of a pair-wise alignment serves as a reference-free metric; we generalize it to the MStA case by averaging over all pairs. Naively, the TM score (Zhang and Skolnick, 2005) could similarly be used; however, an alignment does not suffice to calculate TM because parameters for a rigid body transformation are also required. LDDT (Mariani et al., 2013) is a pair-wise score for comparing a predicted structure (model) against a reference structure. In its original formulation, LDDT is not symmetrical (the calculation is performed from the model’s perspective), and requires that the structures are identical except for atom coordinates. Foldmason defines a variant (LDDT-fm in our notation) for the MStA case. Here, we propose a different extension of LDDT to MStA which we call LDDT-mu. In the pair-wise case, LDDT-mu considers a residue only if it is aligned to a residue in the other structure. To achieve symmetry so that LDDT-mu is unchanged if alignment rows are permuted, distances are included if they are *≤ R*_0_ in either structure, where *R*_0_ is the inclusion radius which is set to 15 Angstroms following the original definition of LDDT. For an MStA, LDDT-mu is averaged over all pairs of structures by default. To avoid prohibitively long runtimes on Balifam3D-1k and Balifam3D-10k, LDDT-mu was measured on a random subset of 100 structures from each alignment. Considering several distinctly different metrics enables a check on whether rankings are sensitive to the choice of metric, giving a more balanced picture of algorithm quality.

### Scaling test

Scaling, i.e. memory and execution time as a function of input data size, was assessed for MStA software only, as follows. Four sets were selected from Balifam3D-100 at random (PF00009, PF00018, PF00037 and PF00046). From each set, subsets of size *N* = 2, 4, 8, 16, 32 and 64 structures were selected at random. For each algorithm, the maximum memory use and maximum execution time across the four sets was measured for each value of *N*. For algorithms supporting multi-threading, this test was repeated with 1 and 16 threads. All tests were performed on a 20-core Linux server (Intel i9-10900K 3.7GHz CPU with 128 Gb RAM).

### Tested software

Given the large number of published MSA and MStA algorithms, a comprehensive assessment was not practical. We therefore selected MSA and MStA software which has been reported to be accurate and/or scalable, representing the state of the art in our best judgment. Software and versions are summarized in Table 1. UPP2 was not evaluated on Homstrad because it failed to align some sets.

**Table 1.** Tested software. *Name* is the abbreviated name used in figures and tables. *Type* is *aa* for algorithms considering only amino acid sequences, *3D* for algorithms considering only *C_α_* coordinates, and *3D+aa* for algorithms which consider both. UPP2 is described in (Park et al., 2023), it was included to represent a recent *aa* algorithm designed to scale. 01252d9 is the commit hash for Muscle. Foldmason (refined) uses 1,000 refinement iterations (`--refine-iters 1000` option).

| Name | Software | Type | Version |
| --- | --- | --- | --- |
| clustalo | Clustal-omega | aa | v1.2.4 |
| foldmason | Foldmason | 3D+aa | v1-763a428 |
| foldmason-i | Foldmason (refined) | 3D+aa | v1-763a428 |
| mafft | Mafft | aa | v7.526 |
| matt | Matt | 3D | v1.0 |
| muscle-3d | Muscle | 3D+aa | 01252d9 |
| muscle-aa | Muscle | aa | 01252d9 |
| mustang | Mustang | 3D | v3.2.4 |
| upp2 | UPP2 | aa | v4.5.1 |
| usalign | US-align | 3D | v20220924 |

### Contact map

The contact map of a protein is a symmetrical matrix giving all-by-all pair-wise Euclidean distances between its *C_α_* atoms. This matrix is often visualized as a “heat map” (Fig. 3), which enables visual identification of domain architecture, secondary structure and contact clusters (Vehlow et al., 2011). The contact map provides a useful framework for thinking about protein structure alignment. Alignment can be formulated as aligning *C_α_* distance matrices, i.e. contact maps, as pioneered by DALI (Holm and Sander, 1993) for the pair-wise case. Also, LDDT can be viewed as quantifying the amount of difference between two aligned contact maps.

**Figure 1.**
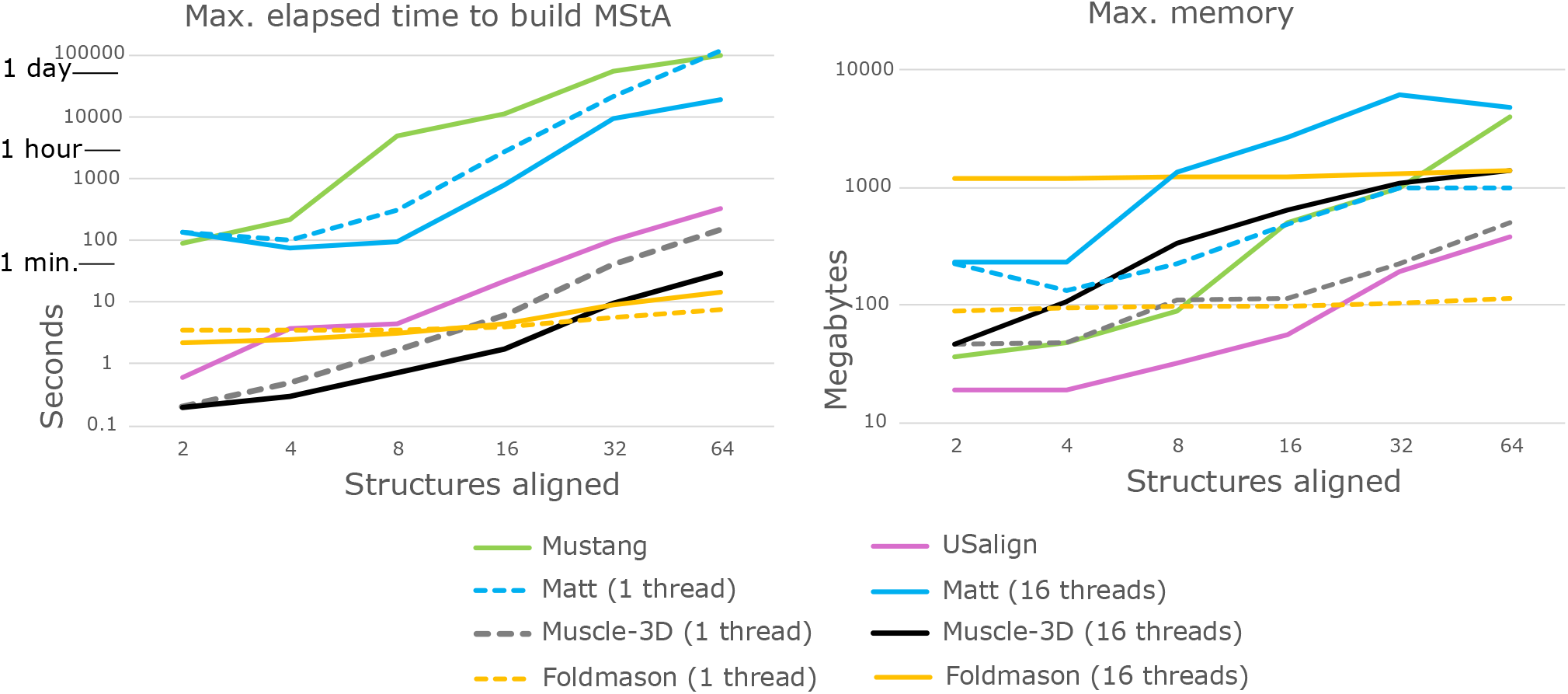
Scaling test results for structure aligners. Maximum elapsed time and memory use for alignments of from 2 to 64 structures (see main text). Note the logarithmic scales.

**Figure 2.**
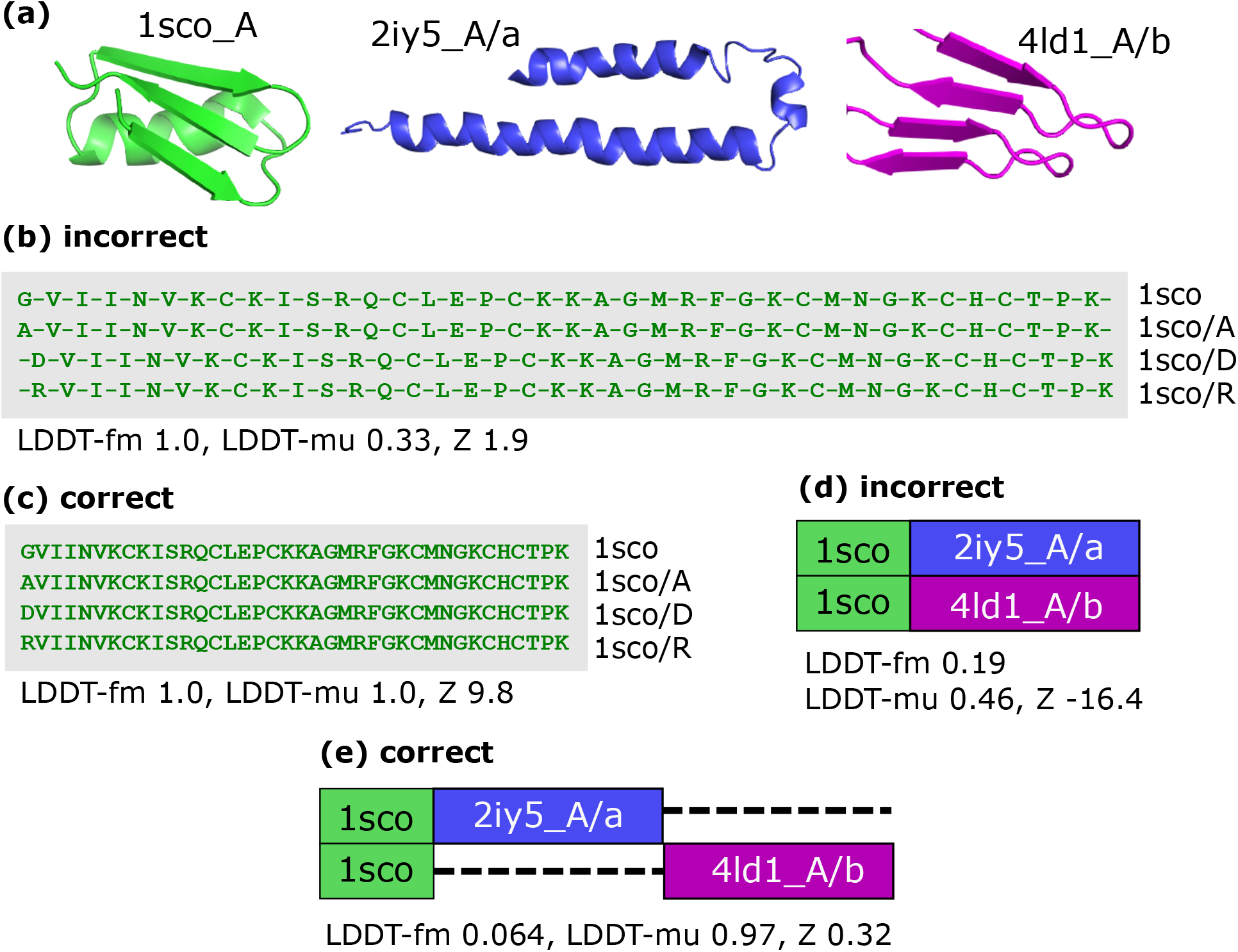
Anomalous accounting for gaps with LDDT-fm. **(a)** Three structures used to construct test cases: (i) scorpion toxin 1sco chain A, (ii) 2iy5 A/a an all-alpha domain at the start of 2iy5 chain A, and (iii) 4dl1 A/b, an all-beta domain at the start of 4ld1 chain A. **(b)** Incorrect alignment of four copies of 1sco A where the first amino acid is changed to ensure that sequences are distinct, e.g. 1sco/D has first letter D. **(c)** Correct alignment corresponding to **(b)**. **(d)** 1sco A+2iy5 A/a aligned to 1sco A+4ld1 A/b; these structure pairs were manually concatenated to simulate multi-domain proteins with one common domain. Domains 2iy5 A/a and 4ld1 A/b are not homologous and have no sequence or structural similarity, therefore **(d)** is incorrect; these domains should be considered independent inserts per alignment **(e)**. Note that LDDT-fm assigns 1.0, the highest possible score, to both **(b)** and **(c)**; also, LDDT-fm assigns a very low score to correct alignment **(e)** and a higher score to incorrect alignment **(d)**. LDDT-mu and *Z* assign higher scores to the correct alignments.

**Figure 3.**
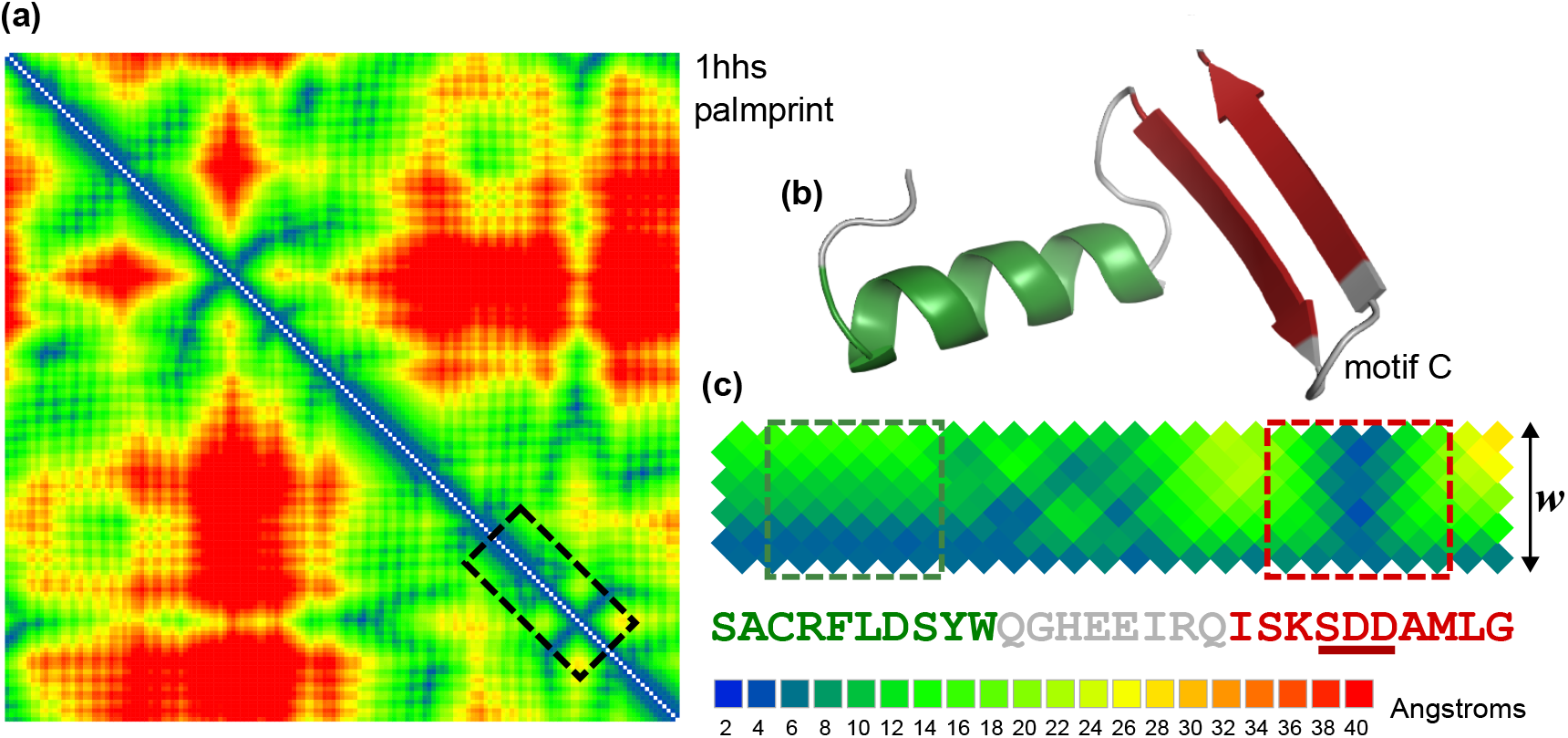
Contact map for 1hhs palmprint. Structure 1hhs is RdRp of bacteriophage Q*β* RdRp, the palmprint is the region spanning from motif A to motif C. A contact map is an all-vs-all distance matrix for *C_α_* atoms which is typically visualized as a “heat map” with colors ranging from green (close) to red (far) as shown in panel (a). The contact map is symmetrical across the diagonal; distance from the diagonal reflects distance in primary sequence while colors reflect distance in 3D space. Panel (b) shows the structure of the segment surrounded by a dashed box in panel (a), this segment comprises an alpha helix followed by motif C, which has a characteristic beta hairpin conformation. Panel (c) shows the band of width *w* = 5 from the dashed box in (a), enlarged and showing only the upper triangle. With small values of *w*, a *w×w* distance matrix centered on a residue reflects its secondary structure, i.e. its local conformation, as illustrated by the dashed boxes in (c).

### Contact map profile

An MStA implicitly aligns several contact maps, enabling analysis of variation in pair-wise *C_α_* atom distances (Iyer et al., 2020). Low variation presumably correlates with evolutionary constraint, while high variation conversely suggests lack of constraint. A natural first step is to calculate the mean and standard deviation of each pair-wise distance, as shown in panels (a) and (b) of Fig. 3. Residue positions now correspond to MStA columns, where “gappy” columns, say with more than 50% gaps, are excluded. Intuitively, variation of, say, *±*2 Angstroms is larger for a pair of atoms at a mean distance of 4 Angstroms than for a pair at 40 Angstroms, leading to the question of how much variation is typically expected for a pair of *C_α_* atoms at a given distance. We addressed this by considering the collection of Balibase reference alignments. Distances in Angstroms were rounded to the nearest integer, and *V* (*d*), i.e. the mean standard deviation for each integer-binned distance *d*, was recorded for each MStA in the collection. The distributions of *V* (*d*) over MStAs was then used to determine whether the standard deviation in distance found for a given pair of positions in a contact map is above or below the typically expected value.

### Local conformation and LDDT-muw

A small sub-matrix of a contact map centered on a residue characterizes its local conformation (Kabsch, 1976), as seen in Fig. 3 (c). In particular, residues in a well-formed alpha helix or beta sheet always have very similar sub-matrices, and a turn is also readily recognized. Many other conformations are possible, generically called loops or coils; these do not cluster into highly similar groups across diverse proteins in the same way as helices or beta sheets. Conservation of local conformation can be recognized by similarity of the corresponding sub-matrices, regardless of whether the secondary structure is regular (strand or sheet). To quantify this, we defined a variant of LDDT-mu derived from a band of width *w* residues around the contact map diagonal (LDDT-muw), which is calculated for a given MStA column *C* as follows. For each row in the MStA, residues in positions *{−w, −w* + 1*…w−* 1*, w}* relative to *C* are identified after discarding gaps. For *N* rows, this gives *N ×* 2*w* residues which are now considered to be aligned to each other, and LDDT-mu is calculated for the implied MStA with 2*w* columns. Discarding gaps ensures the local conformation is reflected in each sub-matrix. If a row has a gap in *C*, the closest column with a letter is used. LDDT-muw ranges from zero to one for each MStA column. This approach could be further generalized to consider a band of primary sequence positions between two distances *w*_0_ and *w*_1_ and a range of 3D distances from *R*_1_ to *R*_0_ where *R*_0_ is the exclusion radius as in the original LDDT and *R*_1_ is an inclusion radius *< R*_0_, enabling quantification of conformation conservation at desired ranges of near/far distances in primary sequence and 3D space; we do not consider this extension further here.

## Results

### Scaling test

Scaling results for aligners which consider structure are summarized in Fig. 1, showing that memory and time scale roughly linearly on a log-log plot. Based on this analysis, we were able to estimate resource requirements for each benchmark and determine which analyses were tractable within our budget constraints. Due to their high computational costs, Matt and Mustang were evaluated on a subset of Balibase but not on the other benchmarks. Foldmason with iterative refinement was evaluated only on Balibase and Homstrad due to the high cost of refinement on large datasets. Only Muscle-3D, Muscle-aa, Mafft and Foldmason were found to be tractable on Balifam3D-10k.

### Balibase3D assessment

Results on Balibase3D using *TC*, *Z*, LDDT-mu and LDDT-fm are summarized in Tables 2-5. Each metric supports a ranking of most algorithms with high statistical significance, but the rankings are quite different except for *Z* and LDDT-mu which generally agree. Most striking is the radical disagreement between LDDT-mu and LDDT-fm. For example, US-align ranks first according to LDDT-mu with *P <* 10*^−^*^11^ against other algorithms, while US-align ranks last according to LDDT-fm with *P <* 10*^−^*^7^.

**Table 2.** Method comparison on Balibase3D. Methods are sorted by decreasing *TC*. Columns with method names report Wilcoxon rank test comparisons with *P* values. “ *>* ” indicates that the method in the row is better than the method in the column with the given *P*-value, similarly “ *<* ” indicates that the row is worse than the column. If *P >* 0.05, “^~^” indicates that the relative ranking is unresolved.

|  | TC | muscle-3d | muscle-aa | clustalo | upp2 | foldmason | foldmason-i | usalign | mafft |
| --- | --- | --- | --- | --- | --- | --- | --- | --- | --- |
| muscle-3d | 0.6522 |  | >5.2e-05 | >2.1e-10 | >9.1e-16 | >4.5e-20 | >2.1e-20 | >4.9e-14 | >2.3e-20 |
| muscle-aa | 0.6060 | <5.2e-05 |  | >0.00011 | >3e-11 | >5.1e-08 | >9.7e-09 | >2.7e-05 | >1.1e-18 |
| clustalo | 0.5770 | <2.1e-10 | <0.00011 |  | >0.0056 | >0.0036 | >0.00053 | ~0.085 | >5.4e-08 |
| upp2 | 0.5492 | <9.1e-16 | <3e-11 | <0.0056 |  | ~0.075 | >0.012 | ~0.73 | >0.00014 |
| foldmason | 0.5328 | <4.5e-20 | <5.1e-08 | <0.0036 | ~0.075 |  | >1.2e-06 | ~0.51 | ~0.085 |
| foldmason-i | 0.5255 | <2.1e-20 | <9.7e-09 | <0.00053 | <0.012 | <1.2e-06 |  | ~0.33 | ~0.5 |
| usalign | 0.5113 | <4.9e-14 | <2.7e-05 | ~0.085 | ~0.73 | ~0.51 | ~0.33 |  | ~0.2 |
| mafft | 0.5081 | <2.3e-20 | <1.1e-18 | <5.4e-08 | <0.00014 | ~0.085 | ~0.5 | ~0.2 |  |

**Table 3.**
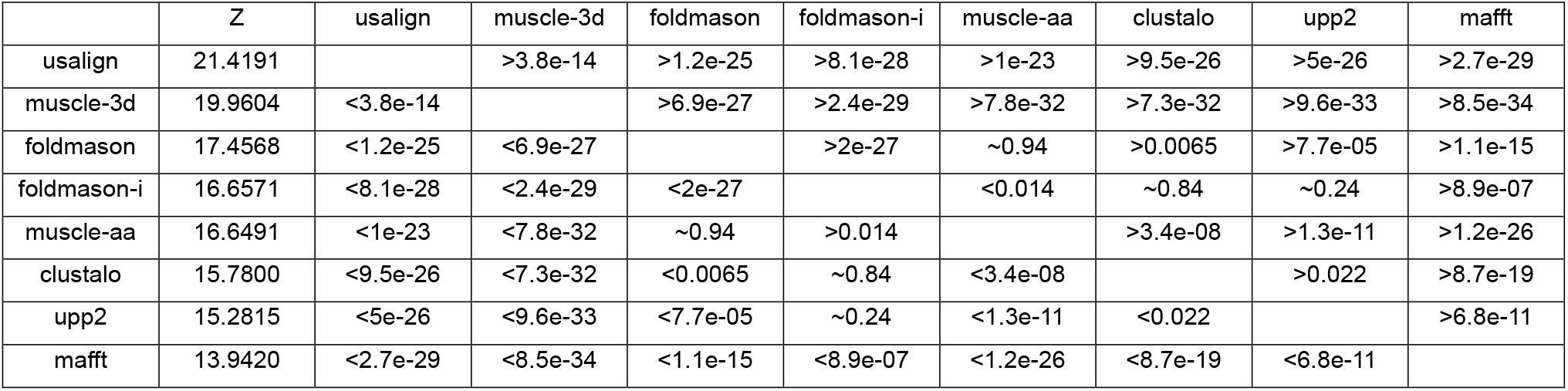
Method comparison on Balibase3D. As Table 2 except that methods are ranked by *Z*.

**Table 4.**
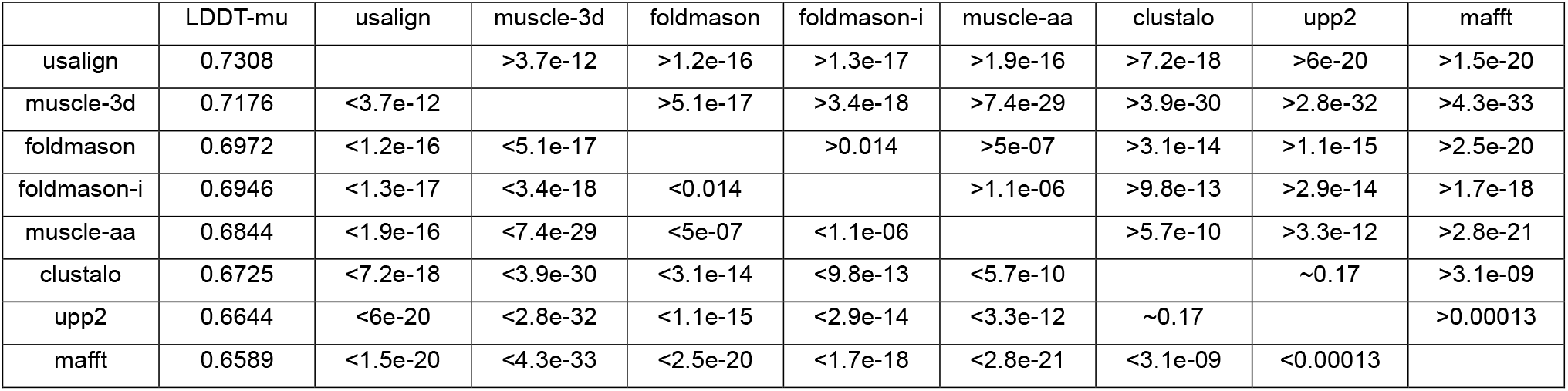
Method comparison on Balibase3D. As Table 2 except that methods are ranked by LDDT-mu.

**Table 5.** Method comparison on Balibase3D. As Table 2 except that methods are ranked by LDDT-fm.

|  | LDDT-fm | foldmason-i | foldmason | clustalo | muscle-3d | upp2 | mafft | muscle-aa | usalign |
| --- | --- | --- | --- | --- | --- | --- | --- | --- | --- |
| foldmason-i | 0.5074 |  | >2e-33 | >4.9e-23 | >2.1e-27 | >1.9e-33 | >1.2e-33 | >1.4e-33 | >3e-30 |
| foldmason | 0.4840 | <2e-33 |  | >0.00034 | >8.9e-12 | >4.6e-22 | >1.5e-26 | >2.4e-28 | >6e-25 |
| clustalo | 0.4692 | <4.9e-23 | <0.00034 |  | >0.00015 | >1.2e-20 | >3.1e-29 | >1.8e-25 | >1.7e-17 |
| muscle-3d | 0.4540 | <2.1e-27 | <8.9e-12 | <0.00015 |  | >3.4e-06 | >6.4e-09 | >4.9e-18 | >4.3e-17 |
| upp2 | 0.4362 | <1.9e-33 | <4.6e-22 | <1.2e-20 | <3.4e-06 |  | >0.0096 | >0.0026 | >9.2e-10 |
| mafft | 0.4290 | <1.2e-33 | <1.5e-26 | <3.1e-29 | <6.4e-09 | <0.0096 |  | ~0.18 | >5.6e-08 |
| muscle-aa | 0.4248 | <1.4e-33 | <2.4e-28 | <1.8e-25 | <4.9e-18 | <0.0026 | ~0.18 |  | >4.9e-07 |
| usalign | 0.3797 | <3e-30 | <6e-25 | <1.7e-17 | <4.3e-17 | <9.2e-10 | <5.6e-08 | <4.9e-07 |  |

### Anomalous accounting for gaps in LDDT-fm

The discrepancy between LDDT-mu and LDDT-fm prompted us to investigate LDDT-fm more closely. The design of LDDT-fm suggested to us that its accounting for gaps could produce undesirable results. We verified this on hand-constructed test cases as shown in Fig. 2. Panel **(b)** shows an alignment of four structures which are identical except for the first amino acid to ensure that sequences are distinct. Gaps were manually introduced into the MStA such that many residue pairs are misaligned by one position; the correct alignment is shown in panel **(c)**. LDDT-fm assigns 1.0, the highest possible score, to both the incorrect MStA **(b)** and the correct alignment **(c)**. By contrast, LDDT-mu is 1.0 for the correct alignment and 0.33 for the misaligned case. The *Z* score is also higher for the correct alignment. Panels **(d)** and **(e)** show alignments of two manually concatenated structures where an identical common domain is followed by two very different domains (all-alpha and all-beta, respectively) to simulate a multi-domain protein with one common domain. The correct alignment is shown in panel **(e)** where offsetting gaps are introduced to indicate that the second domains should be considered independent insertions. Panel **(d)** shows an incorrect alignment where the unrelated domains are aligned to each other. LDDT-fm gives a very low score (0.064) to the correct alignment and a higher score (0.19) to the incorrect alignment. LDDT-mu gives an almost perfect score (0.97) to the correct alignment and low score (0.46) to the incorrect alignment. *Z* also assigns the correct alignment a higher score. These results show that in **(b)** incorrect gaps are not penalized by LDDT-fm, and in **(e)** required gaps are penalized. These issues are avoided by LDDT-mu which averages over pairs of MStA rows rather than over columns. Therefore, we focus primarily on *TC*, *Z* and LDDT-mu for quality assessment. Data and code to reproduce these examples is deposited at https://github.com/rcedgar/pylddt.

### Balifam3D assessment

Results on Balifam3D are shown in Table 6. Rankings by *Z* and LDDT-mu agree, as on Balibase3D. These results show that for larger numbers of structures, including structural information does not necessarily give better results on a structure-based metric as Foldmason consistently has lower LDDT- mu compared to the a.a. methods Muscle-aa and Mafft.

**Table 6.** Method comparison on Balifam3D. *Z* is *Z* score, *mu* is LDDT-mu, (100) is Balifam3D-100, (1k) is Balifam3D-1k, (10k) is Balifam3D-10k. Method rankings according to *Z* and LDDT-mu agree, and are statistically significant except for Balifam3D-10k which has only four sets, and Mafft and Foldmason are not distinguished by any metric.

|  | Z(100) | Z(1k) | Z(10k) | mu(100) | mu(1k) | mu(10k) |
| --- | --- | --- | --- | --- | --- | --- |
| muscle-3d | 12.9 | 9.2 | 2.5 | 0.69 | 0.67 | 0.66 |
| muscle-aa | 10.7 | 7.1 | 1.2 | 0.66 | 0.63 | 0.59 |
| mafft | 8.5 | 4.2 | -1.4 | 0.63 | 0.58 | 0.50 |
| foldmason | 8.5 | 3.8 | -1.1 | 0.61 | 0.51 | 0.32 |

### Homstrad assessment

Results on Homstrad are shown in Table 7. We regard Homstrad as a less stringent test given that most alignments are pair-wise and all proteins have the same domain organization. Method rankings are seen to vary with choice of metric and also if measured on cases with four or more structures.

**Table 7.** Method comparison on Homstrad. Metrics *TC*, *Z* and LDDT-mu on Homstrad. “4+” are metrics measured on sets with at least four structures, noting that Homstrad alignments are mostly pair-wise. Methods are sorted by decreasing *TC*4+.

|  | TC4+ | TC | Z4+ | Z | LDDT-mu4+ | LDDT-mu |
| --- | --- | --- | --- | --- | --- | --- |
| mustang | 0.76 | 0.86 | 20.57 | 21.56 | 0.77 | 0.78 |
| usalign | 0.74 | 0.87 | 21.40 | 22.79 | 0.78 | 0.79 |
| muscle-3d | 0.74 | 0.86 | 18.74 | 19.95 | 0.73 | 0.75 |
| foldmason | 0.71 | 0.86 | 18.22 | 21.43 | 0.74 | 0.77 |
| foldmason-i | 0.70 | 0.86 | 17.98 | 21.35 | 0.74 | 0.77 |
| muscle-aa | 0.69 | 0.80 | 16.27 | 16.22 | 0.71 | 0.71 |
| mafft | 0.67 | 0.79 | 15.64 | 15.48 | 0.71 | 0.71 |
| clustalo | 0.66 | 0.79 | 15.56 | 15.89 | 0.70 | 0.71 |

### Matt and Mustang assessments

Due to their high computational costs, Matt and Mustang were assessed only on smaller alignments in Balibase3D; 89 for Matt and 106 for Mustang. Results are in Supplementary Tables. Both methods were among the highest scoring, but as with other benchmark tests rankings are inconsistent according to different metrics.

### Expected contact map variation

The distribution of *V* (*d*) derived from Balibase is shown in Fig. 7. This distribution was divided into quintiles for each *d*, i.e. four thresholds dividing the distribution into five groups with equal frequencies. This procedure defines categories which we call highest, high, neutral, low and lowest constraint, respectively, enabling plots of constraint as shown in Fiġreffig:profiles.

**Figure 4.**
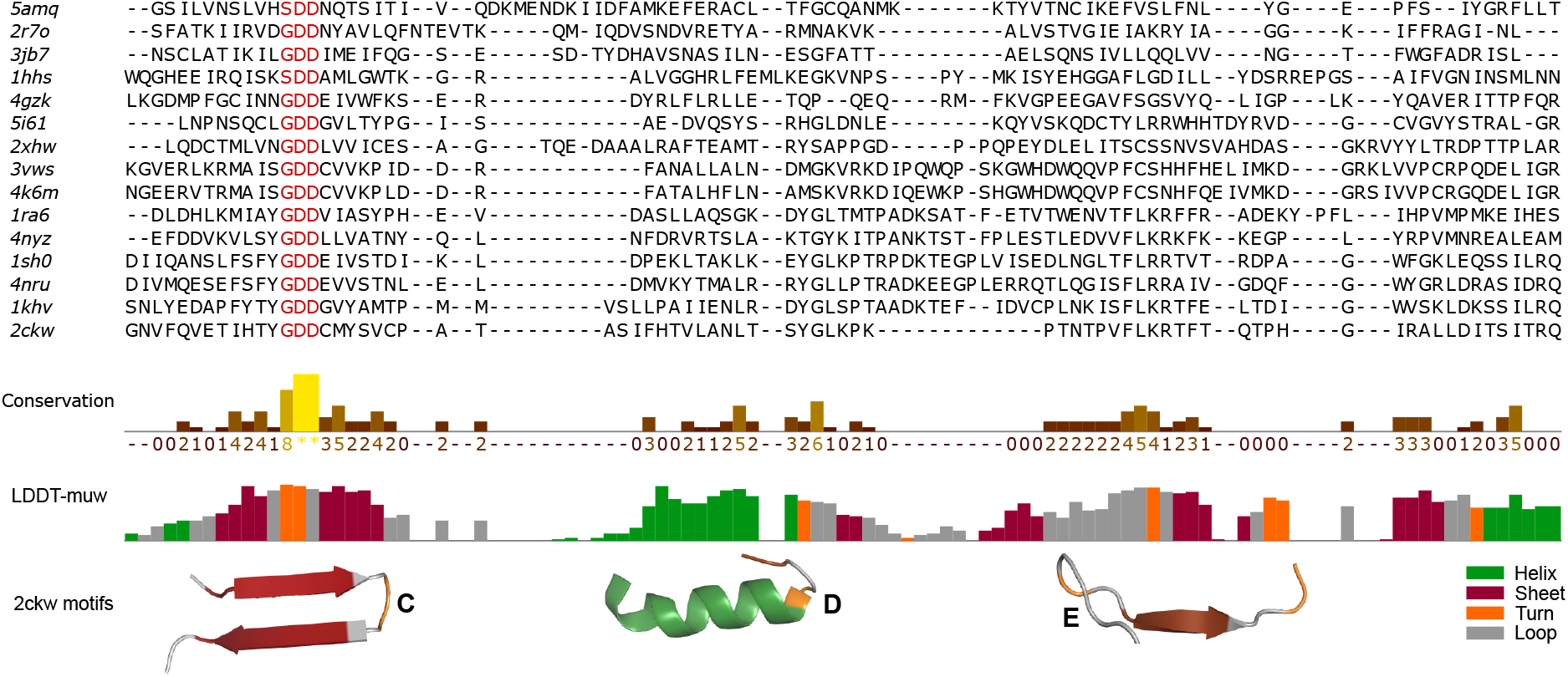
Jalview annotation of LDDT-muw. Muscle-3D alignment of selected solved viral RdRp structures is shown in Jalview showing the segment from motif C to motif E. Annotations shown below the sequence alignment are Conservation, a standard Jalview annotation, together with the LDDT-muw annotation generated by Muscle-3D. LDDT-muw histogram bars are automatically colored according to a four-state secondary structure assignment *{*helix, sheet, turn, loop*}*. Structure cartoons from Pymol were manually added below the LDDT-muw histogram. The well-conserved beta hairpin of motif C is reflected in the sheet-turn-sheet coloring of LDDT-muw, where the turn is placed at the `GDD` sequence motif. Motif D comprises an alpha helix followed by a short loop which is exposed into the catalytic site. Motif E has a short exposed loop followed by a buried beta sheet. Unlike motif C, motifs D and E have minimal sequence conservation and are apparent in distantly-related viruses only when structural conservation is considered.

**Figure 5.**
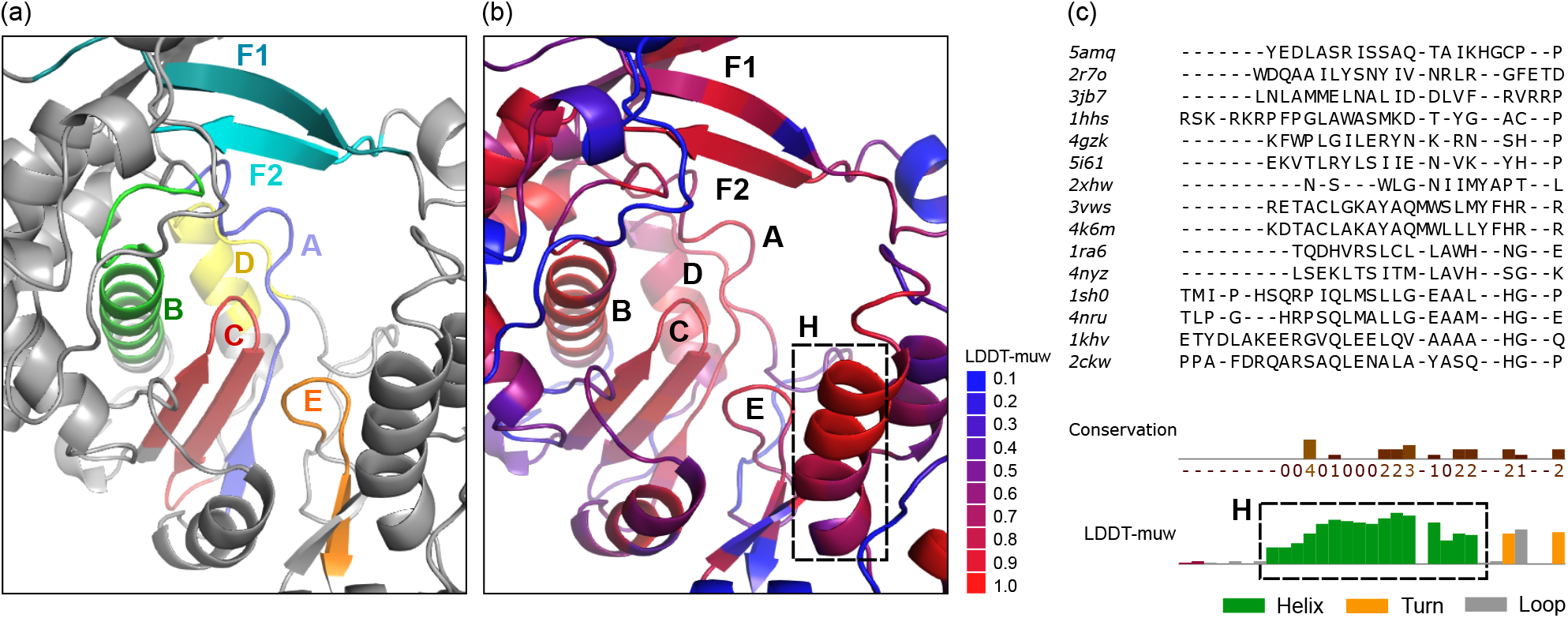
Structure conservation analysis of 1ra6 with Pymol and Jalview. PDB 1ra6 is human poliovirus RdRp. (a) Pymol view colored automatically by Palmscan-3D (https://github.com/rcedgar/palmscan2) to show the canonical structurally-conserved motifs A, B, C, D, E, F1 and F2. (b) Similar Pymol orientation colored automatically by Muscle-3D to show conservation of local conformation according to LDDT-muw. (c) Jalview alignment with LDDT-muw annotations generated by Muscle-3D. Note that high LDDT-muw correlates well with the canonical motifs. A well-conserved helix (H) not included in the canonical motifs is apparent in (b) and (c).

**Figure 6.**
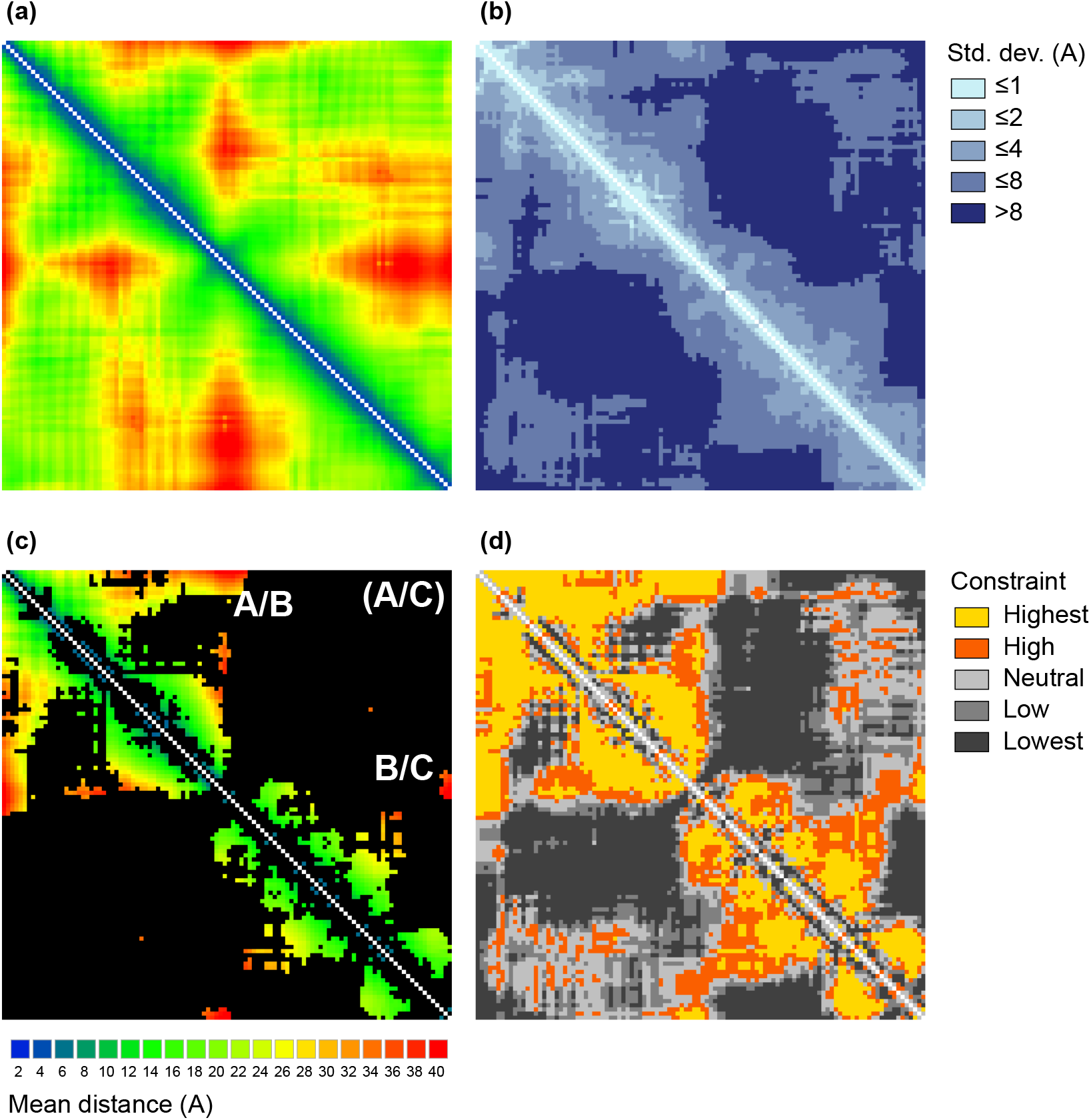
Contact map profiles. (a) Consensus contact map for RdRp palmprint, giving mean all-vs-all distances between *C_α_* atoms in MStA columns with few gaps. (b) Distance conservation measured as standard deviations. (c) Pairs with highest constraint (see panel (d)) colored by distance. (d) Distance constraints in five bins (see main text). High steric constraints between residues at long distances are seen as orange and red regions in panel (c). This analysis shows that motifs A and B have strong long-distance constraints, also motifs B and C, while motifs A and C do not (explained because A and C are spatially close, as shown by green colors in the top-right corner of panel (a)).

**Figure 7.**
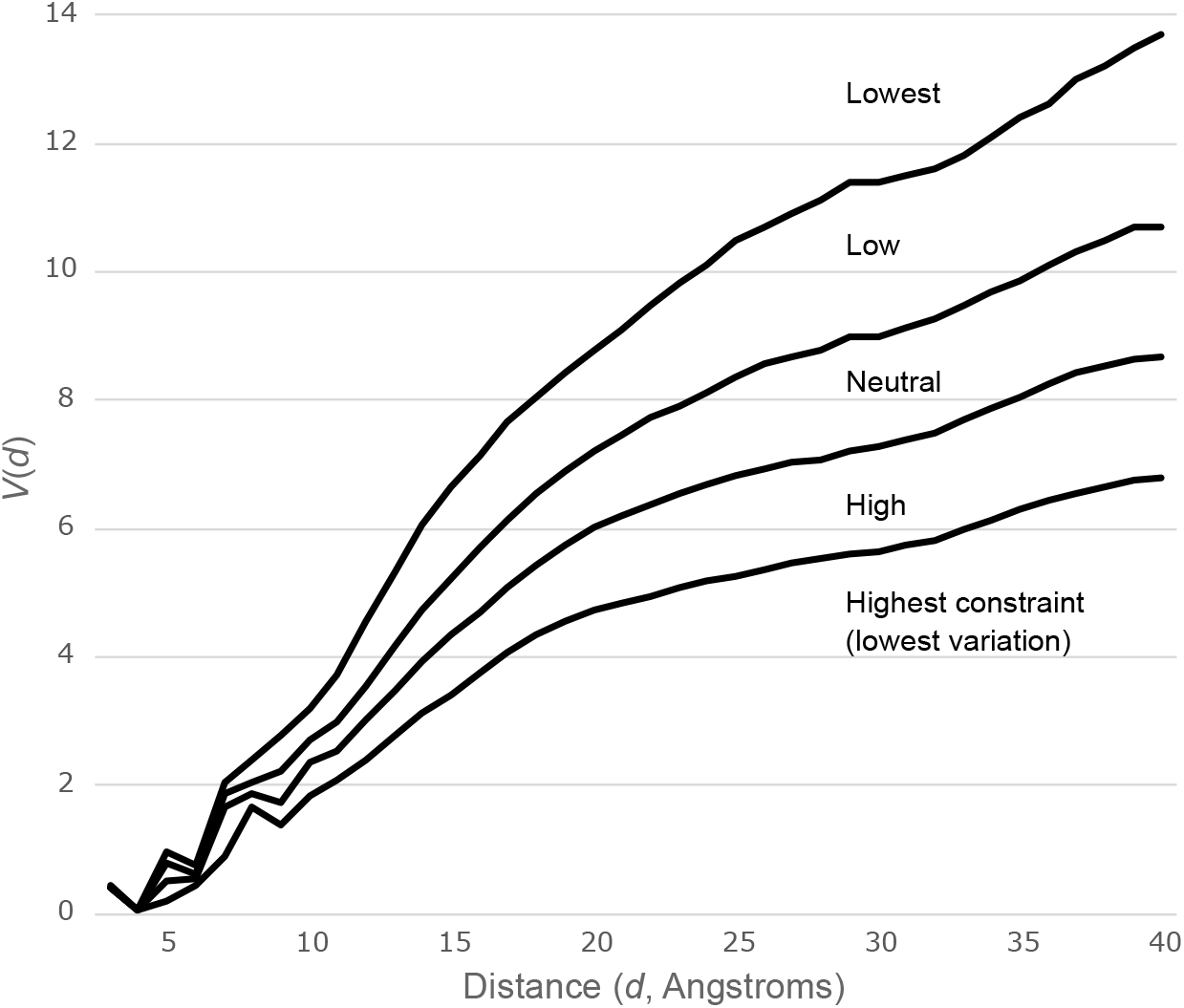
Distribution of *V* (*d*) on Balibase. *V* (*d*) is the standard deviation observed for integer distance *d*. The distribution of *V* (*d*) over a diverse collection of MStAs is used to determine whether a value in a particular contact map profile is higher or lower than expected. For example, at distance *d* = 20 Angstroms, the mean *V* (*d*) over Balibase is 6.6. If smaller variation, say *V* = 4, is found for a *C_α_* pair with mean *d* = 20 in a contact map profile, this indicates that the residue distance is more strongly constrained, and conversely larger variation indicates lower constraint. The distribution is divided into quintiles for each *d* bin, giving the four thresholds shown in this figure. These thresholds are used to assign observed *V* (*d*) into five categories where low variation implies high constraint. An example constraint plot is shown in Fig. 6 (d).

### Jalview and Pymol integration

Jalview (Waterhouse et al., 2009) and Pymol (DeLano et al., 2002) are widely-used software packages for viewing and editing MSAs and structures, respectively. Muscle-3D integrates with these tools by enabling visualization of LDDT-muw, as seen in Figs. 4 and 5. For Jalview, LDDT-muw is shown as a histogram, with taller bars indicating higher secondary similarity. Bars are colored according to a four- state secondary structure alphabet, by default helix=green, strand=maroon, turn-orange and loop=gray. For Pymol, residues are colored according to palette of eleven colors for LDDT-muw = 0, 0.1 0.9, 1.0.

By default, redder indicates higher conservation and bluer indicates lower conservation. Fig. 4 shows an alignment of solved RdRp structures in the region from motif C through motif E, showing that motifs D and E have very low sequence similarity and are apparent only by conservation of local, while motif C is identifiable by the well-known `GDD` sequence motif as well as its characteristic beta hairpin. Fig. 5 shows LDDT-muw coloring in Pymol for human poliovirus RdRp by comparison with annotations of the motifs A through F1/2, showing that high LDDT-muw correlates well with the canonical motifs. In addition, Fig. 5 shows that a conserved helix (which we call H) is apparent in both the Pymol and Jalview visualizations but not accounted for in the standard motifs.

## Discussion

You know a good structural alignment when you see one. Secondary structure elements line up, and sequence similarity is consistent with groups that are well-known to substitute such as *{*`DE`*}* and *{*`ILV`*}*. This intuitive assessment is based on two for criteria placing letters in the same MStA column: (1) residue structural role (e.g., second residue in the third helix), and (2) residue homology. Residue ho- mology, i.e. descent from a common ancestral residue in an unbroken lineage, is well-defined in princi- ple, but is unverifiable in most cases and tends to disappear over time, despite conservation of secondary structure, because every deletion drives a residue lineage to extinction. Structural equivalence is inher- ently fuzzy (see Protein of Theseus), and may diverge from residue homology (Cline et al., 2002; Edgar, 2010). To reconcile these potentially conflicting and/or fuzzy criteria, designers of MStA algorithms and benchmarks must distill the objective of MStA into a single metric, i.e. a number which attempts to capture the desired intuition so that larger numbers indicate better alignments. An algorithm attempts to maximize a metric, where it is typically called something like an alignment score, and a benchmark asks which software produces higher metric values. Different metric designs are possible, e.g. sequence similarity might or might not be considered, and different metrics may better reflect the requirements of different downstream applications. For example, if the MStA will be used for phylogenetic tree estima- tion, then residue homology should be the desired objective rather than structural equivalence, because most tree estimation algorithms are based on substitution models.

## Conclusion

In this work, we found that different metrics are inconsistent to a surprising degree, as found in a pre- vious study of pair-wise structural alignment (Kolodny et al., 2005). Purely structural metrics consid- ering only *C_α_* atom coordinates give conflicting results, even when based on the same pair-wise metric (LDDT-fm and LDDT-mu). Structurally-informed, manually-curated references from Balibase and Homstrad also give conflicting rankings. It is not even clear that considering structure necessarily gives better alignments over amino acid sequences alone; for example, Muscle-aa is better than Foldmason according to the a.a.-aware *TC* metric on Balibase (*P* = 4 *×* 10*^−^*^20^) and the purely structural metric LDDT-mu on Balifam-1k (*P* = 0.006). Muscle-3D is highly scalable and often places among the higher- scoring methods on the benchmark tests considered here, but we conclude that general claims regarding alignment accuracy are not supportable for this category of algorithm.

## Author contributions

R.C.E. conceived and directed the study, implemented the scaling test, implemented LDDT and contact map code, analyzed data and wrote the manuscript. I.T. implemented the Mega alphabet in Muscle, implemented Balibase3D and Balifam3D, implemented and performed benchmark tests and edited the manuscript.

## Software and data availability

Muscle is available at https://github.com/rcedgar/muscle. Benchmark analysis code is available at https://github.com/rcedgar/muscle benchmark. Benchmark data is deposited at TODO.

